# Limitations of EBV transformed human Raji B cells as a model for measuring canonical NF-κB activation

**DOI:** 10.64898/2026.07.07.737082

**Authors:** Rachel Kidwell, Christopher D. Scharer

**Affiliations:** Department of Microbiology and Immunology, Emory University School of Medicine, Atlanta, GA 30322

## Abstract

Autoimmune diseases, such as systemic lupus erythematosus (SLE), are underscored by dysregulated B cell function including the production of autoantibodies, skewed population ratios, and aberrant signaling. Given that the family of nuclear factor kappa B (NF-κB) transcription factors govern responses to stimuli, survival, differentiation, and so forth understanding the intricate regulatory network of NF-κB in B cell biology is paramount for unraveling treatments for B cell-linked autoimmune diseases. Here, we focus on a negative regulator of NF-κB signaling, A20 (*TNFAIP3*), that deactivates NF-κB transcription factor translocation through the ubiquitination and deubiquitination of target proteins. Haploinsufficiency in A20 results in an autoimmune phenotype and mutations to A20 have been associated with SLE, suggesting implications to B cell function. To investigate the role of A20 in NF-κB in human B cells, we generated a *TNFAIP3* knockout (KO) Raji cell line. Cells were stimulated with either anti-IgM or Resiquimod (R848) to activate distinct NF-κB signaling pathways. Using qRT-PCR, western blotting, and flow cytometry, we assessed differences in gene expression, protein production, and NF-κB activation. We observed key limitations in using Epstein-Barr virus transformed B cell lines to model inducible NF-κB signaling.

## Introduction

Tumor Necrosis Factor Alpha-Induced Protein 3 (*TNFAIP3)*, also known as A20, was originally identified as a transcriptional response gene in human umbilical vein endothelial cells stimulated with TNF-α, and subsequently characterized as a protective factor limiting TNF-α-induced cell death^1–3^. At homeostasis, A20 is ubiquitously expressed at low levels across multiple cell types; however, its expression is rapidly upregulated following NF-κB pathway activation^4^. The importance of A20 in restraining NF-κB-driven inflammation was determined through the establishment of A20 knockout (A20^-/-^) mice, which developed systemic inflammation and prematurely died one week after birth^5^. These mice also exhibited fatal hypersensitivity to TNF-α and LPS challenges. Collectively, these findings drove research into A20 functional mechanisms and its role as an essential gatekeeper of NF-κB function by maintaining inflammatory homeostasis.

A20 is now recognized as a critical regulator of both the innate and adaptive immune responses, exerting its inhibitory function primarily through control of the ubiquitination machinery that governs NF-κB pathway activation^6,7^. The canonical NF-κB pathway is initiated through diverse stimuli, including TNF, CD40, and BCR signaling, converging on intracellular signaling cascades that activate the NF-κB family of transcription factors and alter target gene expression. Central to this regulation is the post-translational modification of signaling proteins through ubiquitination, whereby ubiquitin (Ub) chains are covalently attached to a substrate lysine residue^8^. The addition of polymeric Ub chains is mediated by a sequential cascade of enzymatic reactions between E1, E2, and E3 ligases. These enzymes transfer Ub to one of seven specific amino acid residues (K6, K11, K27, K29, K33, K48, K63) or the α amino group of the N-terminal methionine (Met1). Depending on the site, the Ub modification results in distinct functional outcomes such as proteasomal degradation, signal transduction, and receptor trafficking^8^. Central to control of the NF-κB pathway is also the process of deubiquitination, in which deubiquitinases (DUBs) remove polyubiquitin chains from substrates^9^.

A20 uniquely controls NF-κB signaling by operating on both sides of the Ub machinery by both adding K48-linked Ub chains to target proteins, tagging them for proteasomal degradation, and removing essential K63-linked chains that are required for productive NF-κB signaling transduction^10,11^. A20 executes these dual Ub-editing functions through two structurally and functionally distinct domains. The N-terminal ovarian tumor binding domain (OTU) (residues 1-370) encodes DUB activity, whereas the C-terminus contains seven repeated zinc finger (ZnF) domains that mediate E3 ligase function and protein-protein interactions^12^. A20 has been documented to interact with several proteins that are key mediators of NF-κB activation, including TRAF2, TRAF6, RIP1, RIP2, and NEMO through these domains^13,14^. The OTU domain is responsible for disassembling the K63 polyUb chains from RIP1 and TRAF2/6, which are required from the recruitment of the NEMO complex and downstream NF-κB activation^11,13^. The catalytic activity of the OTU domain is mediated by the C103 residue, as demonstrated across multiple independent studies^10,13,15,16^. However, the functional necessity of A20 DUB activity has been debated, with some studies indicating it may be dispensable for A20-mediated suppression of inflammation *in vitro*^17,18^. Characterization of a knock-in mouse model harboring a C103A point mutation to eliminate the DUB activity of A20 *in vivo*^19^ did not exhibit aberrant inflammation and lived a normal lifespan. These findings highlight that precise contribution of OTU activity to A20 function remains poorly understood and raises important questions including whether subtle reductions in OTU activity may alter downstream gene expression profiles, whether these findings are translatable to human B cells, and how the OTU domain functionally cooperates with A20’s E3 ligase activity.

Complementing the OTU domain activity, the ZnF motifs mediate A20’s E3 ligase function by adding K48-linked polyUb chains to target proteins, directing them towards proteasomal degradation^13^. ZnF4 has been shown to catalyze K48 Ub of RIP1, promoting its degradation and destabilization of the UBc13 (an E2 ligase) interaction with cIAP1/2, and thus preventing new K63 Ub chains from forming^11^. This finding has been supported by murine models in which deletion of ZnF4 alone leads to increased RIP1 and subsequently increased NF-κB signaling^12,20^. However, ZnF4 deficient mice did not develop uncontrolled inflammation, suggesting that the dual function of A20 may need to be disrupted for lethality in mice.

More recently, both *in vivo* and *in vitro* evidence has suggested ZnF7 is involved in NF-κB pathway shutdown. ZnF7 is thought to inhibit NEMO activation through a nonenzymatic mechanism by binding M1-linked polyUb^21^. Moreover, as another regulatory mode, A20 binds M1-linked polyUb generated by the LUBAC complex, destabilizing the NEMO complex interactions, and attenuating NF-κB activation^22–25^. Consistent with this, mice bearing inactivating mutations in the A20 ZnF7 domain (A20^ZnF7/ZnF7^ knock-in mice) developed a proinflammatory phenotype consisting of high inflammatory cytokine concentrations, splenomegaly, and altered splenic immune cell populations, suggesting ZnF7 regulates immune homeostasis^7^. Together these findings suggest that the OTU domain, ZnF4, and ZnF7 have functionally distinct yet coordinated mechanisms through which A20 suppresses NF-κB activity and maintains immune regulation.

Beyond its well-established role in canonical NF-κB regulation, A20 has been implicated in modulating the transition between the canonical and noncanonical NF-κB^26^. As previously described, the noncanonical NF-κB pathway is activated through NIK following TRAF2/3 and cIAP1/2-mediated K48 ubiquitination and degradation^27–29^. A20-deficient cells demonstrated dysregulated NIK accumulation and reduced p100 processing to p52 following pathway stimulation, resulting in the inability to shift from canonical to noncanonical NF-κB signaling following LTβ stimulation^26^. To our knowledge, this represents the only study examining A20’s role in the molecular switch between both NF-κB pathways, but CD40 signal transduction results in the nuclear localization of both canonical and noncanonical transcription factors to the nucleus, making it plausible there is A20 involvement in pathway crosstalk. This remains an important open question and will require further investigation utilizing various stimuli for the canonical and noncanonical NF-κB pathways.

It is well understood that maintaining B cell homeostasis requires the effective coordination of diverse signaling avenues resulting in NF-κB activation and these are mediated through negative regulation by A20. However, most of the research surrounding A20 has not been conducted in human B cells. Several studies have investigated the role of A20 following TNF-α or lipopolysaccharide (LPS) stimulation in various cell types including human T cells, human embryonic kidney cells (HEK293), epithelial cells, dendritic cells, and murine models^30–35^. This highlights a clear gap in understanding the molecular map of A20 human B cell functioning to regulate NF-κB. The limited number of studies that have examined human B cells have primarily focused on clinical phenotypes associated with *TNFAIP3* haploinsufficiency, including B cell frequencies, serum immunoglobulin (Ig) titers, and proinflammatory cytokine levels instead of the underlying cellular mechanisms of A20 function^36–39^. As a result, there are several remaining questions including how do different stimuli or concurrent stimulation affect A20-mediated NF-κB regulation in human B cells, what is the molecular map of A20 and protein interactions that happen *in vivo* following human B cell stimulation, and how changes in A20 function can result in different NF-κB gene expression patterns. Addressing these questions is central to understanding the contribution of dysregulated A20 function to human B cell biology.

The clinical relevance of A20 is underpinned by the identification of multiple *TNFAIP3* polymorphisms across genome-wide association studies (GWAS)^40^. A20 polymorphisms have been associated with numerous autoimmune diseases such as systemic lupus erythematous (SLE), rheumatoid arthritis (RA), psoriasis, type-1 diabetes, Sjögren’s syndrome, and Crohn’s disease^41^. Multiple *TNFAIP3* polymorphisms, located in both the OTU domain (rs2230926, rs5029927, rs5029941) and ZnF domains (rs5029953 & p.Ala588Valfs*80) have been linked to autoimmune diseases^42–45^. Furthermore, several studies have linked A20 polymorphisms to increased NF-κB signaling in B cell lymphomas^46–48^. Missense mutations at these loci may result in amino acid changes altering the structure and function of the A20 protein. Beyond single nucleotide polymorphisms, GWAS have also revealed frame shift mutations, premature stop codons, missense mutations, and splicing changes that may more completely diminish A20 function. Given A20’s potent anti-inflammatory role in response to NF-κB activation, even modest changes to the protein sequence could have notable consequences on the Ub and DUB functions of A20 across an individual’s lifespan. The association of *TNFAIP3* variants with both autoimmune diseases and cancers further highlight the clinical contexts in which dysregulated A20 function can manifest.

The consequences of A20 loss of function are more strikingly illustrated by haploinsufficiency of A20 (HA20), a Behçet’s-like disease caused by high-penetrance loss-of-function germline mutations in *TNFAIP3*^37,39,49,50^. Affected individuals present with clinical symptoms such as recurrent ulcers, abscesses, and renal or gastrointestinal complications. Functional studies performed on patient-derived cells showed increased degradation of IκBα and nuclear translocation of the NF-κB p65 subunit together with increased expression of NF-κB-mediated proinflammatory cytokines^37,51^. Furthermore, cells harboring the heterozygous germline A20 mutations had defective removal of K63-linked Ub from TRAF6, NEMO, and RIP1 highlighting defective A20 DUB activity^52^. Together, these findings suggest that mutations or insufficiency of A20 may lead to unrestricted NF-κB signaling causing hyperresponsiveness following stimulation and resulting in dysfunctional cells and symptoms of autoimmunity.

The association of A20 polymorphisms and B cell-driven autoimmune diseases, such as SLE and Sjögren syndrome, has motivated researchers to study the effect of A20 polymorphisms in murine B cell models. In contrast to the lethal systemic inflammation observed in global A20-deficent mice, B cell-specific deletion of A20 using a CD19-Cre system, does not cause premature death^53–55^. These mice exhibit an increased number of germinal center B cells, autoantibodies, and autoantibody complexes leading glomerular deposits^54^. Additionally, two independent concurring studies demonstrated that A20 is necessary for MZ B cell formation and noted a hyperactive B cell phenotype leading to enhanced proliferation upon activation leading to splenomegaly^53,55^. These results from murine model studies suggest that loss of A20 function in B cells leads to an autoimmune pathogenesis including an increased production of autoantibodies and uncontrolled inflammatory response. However, these papers did not dissect specific B cell subsets that are now known to be associated with autoimmunity.

In summary, A20 has emerged as an indispensable regulator of NF-κB signaling across immune cell types by the coordination of DUB and Ub activities through its OTU and ZnF domains to prevent aberrant NF-κB activation. Germline mutations, haploinsufficiency, and disease-associated polymorphisms to *TNFAIP3* may result in decreased function leading to unconstrained proinflammatory cytokine release, accumulation of NF-κB transcription factors in the nucleus, and dysregulated gene expression impacting cellular homeostasis. The role of NF-κB in human B cell function is vitally important for B cell maturation, activation, and equilibrium; however, A20 in B cells has remained poorly defined. Establishing how A20 governs NF-κB activation dynamics and gene expression in human B cells is therefore essential to understanding its contribution to B cell-mediated autoimmunity and represents the central focus of this work.

To address these gaps, this study aimed to define the role of A20 in regulating NF-κB signaling and B cell function using complementary experimental approaches. To study A20 in the context of human B cells, we utilized the human B cell line Raji, which was originally derived from a patient with Burkitt lymphoma and was immortalized using Epstein–Barr virus (EBV). We first examined the functional consequences of *TNFAIP3* loss in a Raji B cell knockout model stimulated with anti-IgM and R848 to engage distinct NF-κB signaling pathways, before transitioning to primary human B cells to more accurately model physiologically relevant A20-dependent regulation. Together, these studies seek to characterize how A20 shapes NF-κB activation dynamics, B cell-specific gene expression profiles, and fate decision outcomes, and to establish the broader implications of A20 dysregulation for autoreactive B cell expansion in the context of autoimmunity.

## Results

### CRISPR-mediated knockout of A20 in Raji B cells

To investigate the role of A20 in NF-κB signaling in human B cells, we generated A20-deficient Raji B cells using CRISPR/Cas9-mediated knockout of *TNFAIP3*. Prior to screening, a sequential gating strategy was established to identify lymphocytes, single cells, and live cells before evaluating A20 protein expression by flow cytometry (**Fig. 1A**). Four gRNAs targeting *TNFAIP3* were screened by intracellular flow cytometry to assess A20 protein loss with RNA gRNA #2 and #5 demonstrating the greatest reduction in A20 median fluorescence intensity (MFI) compared to the non-targeting (NT) and isotype controls (**Fig. 1B-C**). gRNA #1 demonstrated no reduction in A20 protein expression and #6 had a modest reduction. Therefore, we continued to confrim parental clones #2 and #5 by western blot to ensure our flow cytometry results were not due to nonspecific binding of the antibody (**Fig. 1D**). Western blot results confirmed a near complete loss of A20 protein expression in cells that received gRNA #2 and #5 compared to wildtype (WT) Raji cells. To generate clonal knockout lines and reduce heterogeneity within the population of knockout (KO) cells, we single-cell sorted the parental clones #2 and #5 using lymphocyte and live markers (**Fig. 1E**). After expansion of the sorted clones, we validated A20 protein reduction through flow cytometry and MFI calculation (**Fig. 1F**). The resulting MFI showed that several of the expanded clones had reduced A20 MFI compared to parental clones #2 and #5, including P2C2, P5C10, C2C15, P5C16, P5C19, and P5C20. Specifically, P2C2 and P5C10 had the lowest MFI and were selected for downstream validation via Sanger sequencing to confirm genomic disruption of the *TNFAIP3* gene.

**Figure 1.**
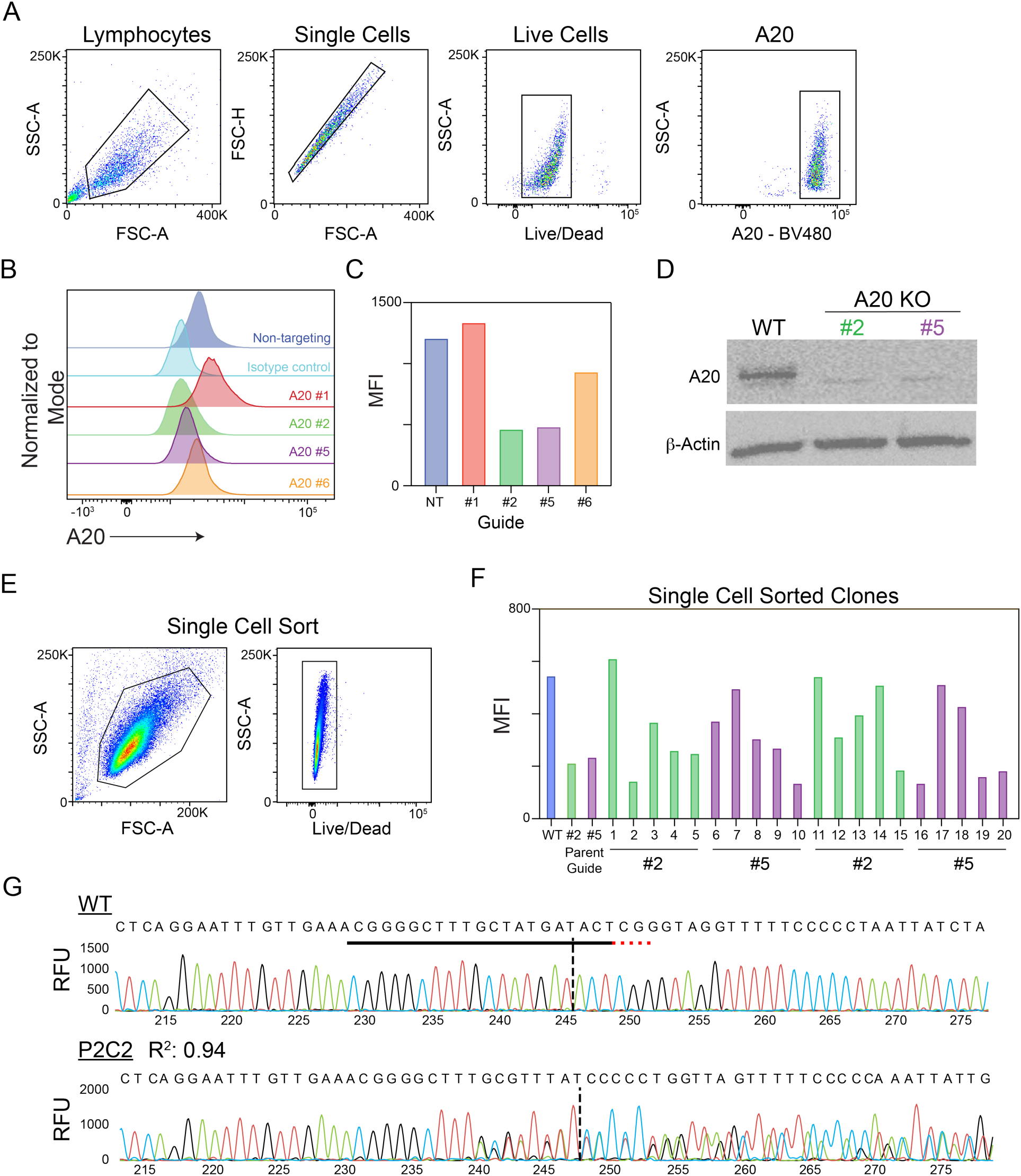
Generation and validation of CRISPR/Cas9-mediated A20 knockout Raji B cells. **A.** Representative sequential flow cytometry gating strategy used to identify lymphocytes (FSC-A vs SSC-A), single cells (FSC-A vs FSC-H), live cells (Live/Dead), and A20-expressing cells (A20-BV480) prior to screening of CRISPR gRNAs. **B.** Flow cytometry histograms showing A20 protein expression in Raji cells transduced with four CRISPR gRNAs targeting TNFAIP3 (A20 #1, #2, #5, #6) compared to non-targeting (NT) and isotype controls. **C.** Quantification of A20 median fluorescence intensity (MFI) across gRNAs and NT control. **D.** Western blot validation of A20 protein loss in parental knockout clones generated with gRNA #2 and #5 compared to wildtype (WT) Raji cells. β-actin serves as a loading control. **E.** Gating strategy used for single cell sorting of parental knockout clones, including lymphocyte (FSC-A vs SSC-A) and live cell (Live/Dead) gates. **F.** A20 MFI of single cell sorted and expanded clones derived from parental gRNA #2 and #5, compared to WT and parental guide controls. **G.** Sanger sequencing traces of the *TNFAIP3* locus in WT and the P2C2 clone. The predicted CRISPR cut site is indicated by the dashed vertical line. The black horizonal line represents the P2C2 guide placement, and the red dots are the protospacer adjacent motif (PAM) site.

To confirm on-target genomic editing, we isolated genomic DNA from P2C2 and P5C10 and performed Sanger sequencing to verify disruption of the *TNFAIP3* locus (**Fig. 1G**). Editing efficiency and indel profiles were analyzed using the Inference of CRISPR Edits (ICE)^56^ algorithm. Analysis of the P2C2 clone demonstrated an insertion-deletion (indel) mutation at the predicted cut site with a high confidence (R^2^ = 0.94), confirming efficient CRISPR-medicated editing of the *TNFAIP3* gene. Based on the depth of protein loss and genomic confirmation, P2C2 was selected as the primary clone for subsequent downstream experiments, with P5C10, P5C19, and P5C20 retained as confirmatory clones where indicated. Together, these findings confirm robust and specific loss of A20 protein expression in CRISPR/Cas9-edited Raji B cells, providing a validated experimental cell line for interrogating A20-dependent NF-κB regulation in human B cells.

### Raji cells demonstrate limited inducible NF-κB signaling in response to R848 stimulation

As a crucial negative regulator of canonical NF-κB signaling, A20 is predicted to limit NF-κB translocation to the nucleus; therefore, loss of A20 may result in enhanced and prolonged nuclear localization. Previous studies have demonstrated that alterations in the A20 protein that reduce its function led to increased NF-κB signaling; however, these studies were not performed in human B cells^57^. Herein, we sought to establish a reliable stimulation system capable of inducing measurable canonical NF-κB activation as a functional readout of A20 loss in human B cells. To assess whether TLR stimulation could activate NF-κB signaling in Raji cells, we utilized the synthetic compound, Resiquimod (R848), a dual agonist of TLR7 and TLR8 in human cells which has been shown to induce NF-κB activation *ex vivo* in monocytes^53^ and potently activate primary human B cells^58,59^. To validate A20 protein expression following stimulation, we performed western blot analysis on WT Raji cells and A20-deficient clones P2C2, P5C19, and P5C20 under both unstimulated and 4-hour (hr) R848 stimulated conditions (**Fig. 2A**). WT Raji cells exhibited robust A20 expression under both unstimulated and stimulated conditions, with a modest increase observed following R848 stimulation. In contrast, A20 protein was markedly absent or greatly reduced (clone P5C20) in A20-deficient clones regardless of stimulation. Quantification of A20 normalized to β-actin further demonstrated a slight increase of A20 in WT Raji cells following stimulation and confirmed the negligible A20 expression in the knockout clones before and after stimulation (**Fig. 2B**). Together, these results demonstrate that R848 stimulation induces A20 expression in WT Raji cells, consistent with activation of an NF-κB-dependent negative feedback loop and provide a foundation for assessing NF-κB signaling in these cells.

**Figure 2.**
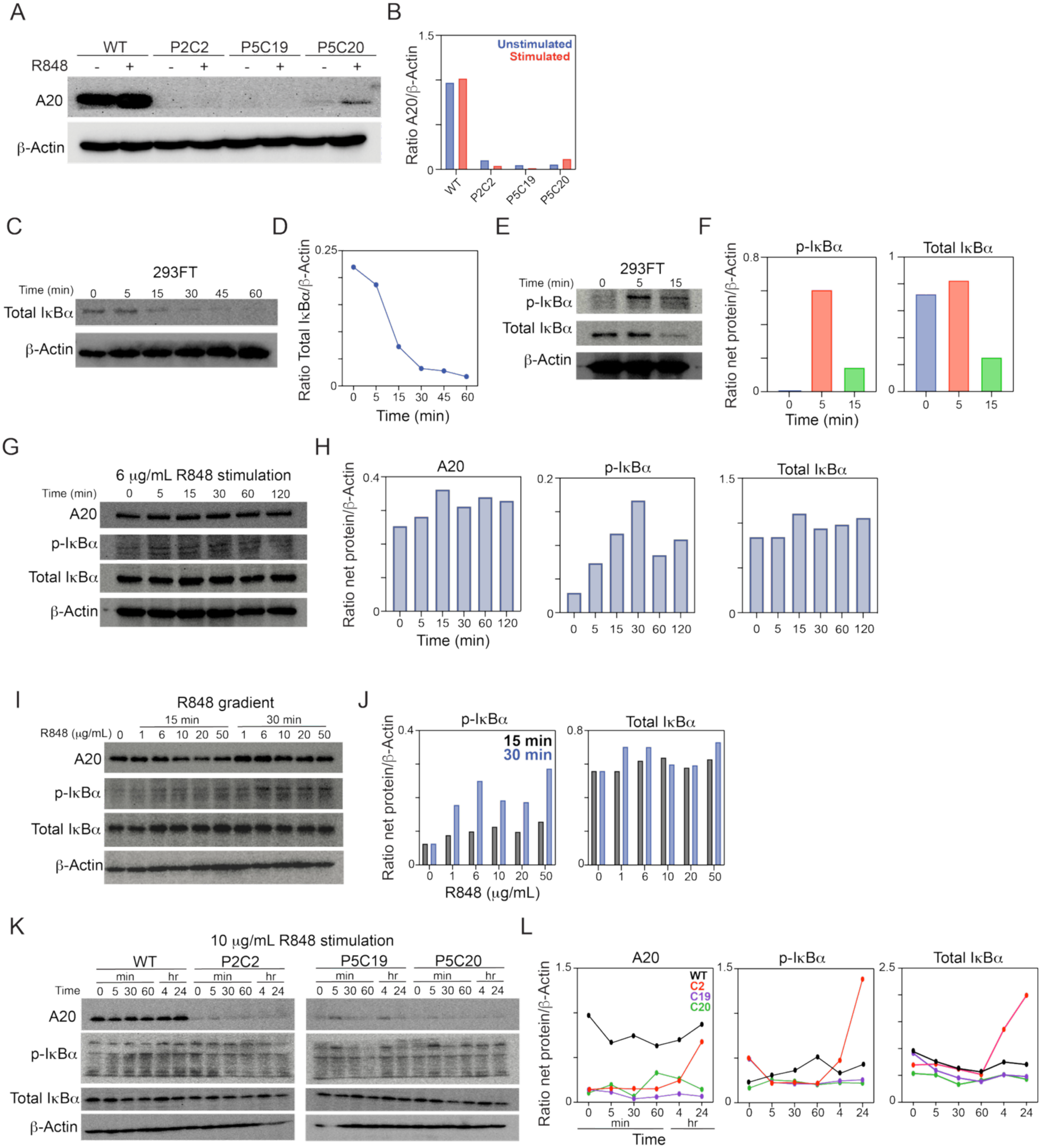
Raji cells demonstrate limited inducible NF-κB signaling in response to R848. **A.** Western blot analysis of A20 protein expression in WT and A20-deficient Raji clones (P2C2, P5C19, P5C20) under unstimulated (-) and R848-stimulated (+) conditions. β-actin serves as a loading control. **B.** Quantification of A20 expression normalized to β-actin in unstimulated (blue) and R848-stimulated (red) conditions across WT and A20-deficient clones. **C.** Western blot time course of total IκBα expression in 293FT cells following TNF-α stimulation at the indicated time points (0–60 min). β-actin serves as a loading control. **D.** Quantification of total IκBα normalized to β-actin, demonstrating time-dependent IκBα degradation consistent with canonical NF-κB activation kinetics. **E.** Western blot analysis of p-IκBα, total IκBα, and β-actin in 293FT cells at 0, 5, and 15 minutes following TNF-α stimulation. **F.** Quantification of p-IκBα and total IκBα normalized to β-actin from **E**. **G.** Western blot time course of A20, p-IκBα, total IκBα, and β-actin in WT Raji cells stimulated with R848 (6 µg/mL) at indicated time points (0-120 min). **H.** Quantification of A20, p-IκBα, and total IκBα normalized to β-actin from **G**. **I.** Western blot analysis of A20, p-IκBα, total IκBα, and β-actin in WT Raji cells treated with increasing concentrations of R848 (0-50 µg/mL) at 15- and 30-min. **J.** Quantification of p-IκBα and total IκBα normalized to β-actin from **I** at 15 min (blue) and 30 min (grey). **K.** Western blot time course of A20, p-IκBα, and total IκBα in WT and A20-deficient Raji clones (P2C2, P5C19, P5C20) stimulated with R848 (10 µg/mL) across 0–24 hours. β-actin serves as a loading control. **L.** Quantification of A20, p-IκBα, and total IκBα normalized to β-actin from (K) across all cell lines and time points.

To directly assess canonical NF-κB activation through IκBα phosphorylation and degradation, we first validated the specificity and sensitivity of our p-IκBα and total IκBα antibodies using human embryonic kidney (293FT) cells as a well-established in vitro model for NF-κB activation^60,61^. Following stimulation with TNFα, 293FT cells demonstrated a time-dependent degradation of total IκBα that was largely complete by 30 minutes (min), accompanied by a clear increase in p-IκBα at 5 min that subsequently resolved by 15 min, consistent with canonical NF-κB activation kinetics (**Fig. 2C-F**). These results confirm that our antibodies reliably detect IκBα phosphorylation and degradation with appropriate sensitivity and specificity, validating their use for downstream experiments in Raji cells.

To confirm NF-κB pathway activation in Raji cells, we performed a time-course analysis, following stimulation of WT cells with R848 (6 μg/mL) using the same antibodies confirmed in Fig. 2C-F (**Fig. 2G**). In contrast to the clear degradation of total IκBα and p-IκBα in 293FT cells, Raji cells did not exhibit a robust decrease in total IκBα or induction or p-IκBα over the same time course (**Fig. 2H**). We did observe a peak of p-IκBα at 30 min with a striking decrease at 60 min; however, this does not follow previously published data of NF-κB kinetics in 293FT cells stimulated with TNFα^61^. Furthermore, levels of A20 in WT Raji cells peaked at 15 min but did not show significant changes throughout the time course. Additionally, we evaluated whether this response was dependent on stimulus strength, Raji cells were treated with increasing concentrations of R848 and analyzed at 15- or 30-minute time points (**Fig. 2I-J**). However, even at higher concentrations up to 50 μg/mL of R848, we did not observe visible IκBα degradation or increase in p-IκBα levels. Overall, these results highlight a lack of inducible NF-κB activation and suggest constitutive expression of A20, IκBα, and p-IκBα in Raji cells.

Given the lack of NF-κB activation observed in WT Raji cells following R848 stimulation, we next investigated whether removal of the A20 protein would enhance pathway responsiveness. WT and A20-deficient Raji cell clones (P2C2, P5C19, and P5C20) were stimulated with R848 (10 μg/mL) and assessed for IκBα phosphorylation and degradation (**Fig. 2K-M**). In contrast to WT Raji cells, which exhibited minimal changes in p-IκBα and no clear evidence of IκBα degradation, A20-deficient clones demonstrated variable alterations in NF-κB signaling. Specifically, the P2C2 clone exhibited a marked increase in p-IκBα at later time points, whereas other clones showed more modest or limited changes. Despite this, canonical early-phase IκBα degradation kinetics were not observed in either WT or A20-deficient Raji cells. Together, these findings suggest that while a loss of A20 may enhance or prolong NF-κB signaling in certain contexts or cell types, Raji cells do not exhibit inducible NF-κB activation in response to R848 stimulation under varying conditions.

### Limited detection of RelA/p65 nuclear localization in Raji cells

Because of the limited ability to detect inducible NF-κB activation in Raji cells when investigating IκBα phosphorylation and degradation, we next assessed whether nuclear translocation of the canonical NF-κB transcription factor, RelA/p65 could serve as an alternative readout of pathway activation. Previously published studies have successfully isolated nuclei from stimulated immune cells and quantified the expression of RelA/p65 localization to demonstrate NF-κB activation^62^.

To establish this approach in our system, we first optimized nuclear isolation in Raji cells and evaluated isolated nuclei and whole cells by flow cytometry (**Fig. 3A**). Forward and side scatter properties differed between isolated nuclei and intact cells, consistent with successful separation. To further validate nuclear isolation, we assessed the expression of the B cell surface marker CD19 (**Fig. 3B**-**C**). As expected, isolated nuclei exhibited minimal CD19 staining, whereas whole cells were predominantly CD19^+^, confirming the loss of cell surface membrane signal during nuclear isolation.

**Figure 3.**
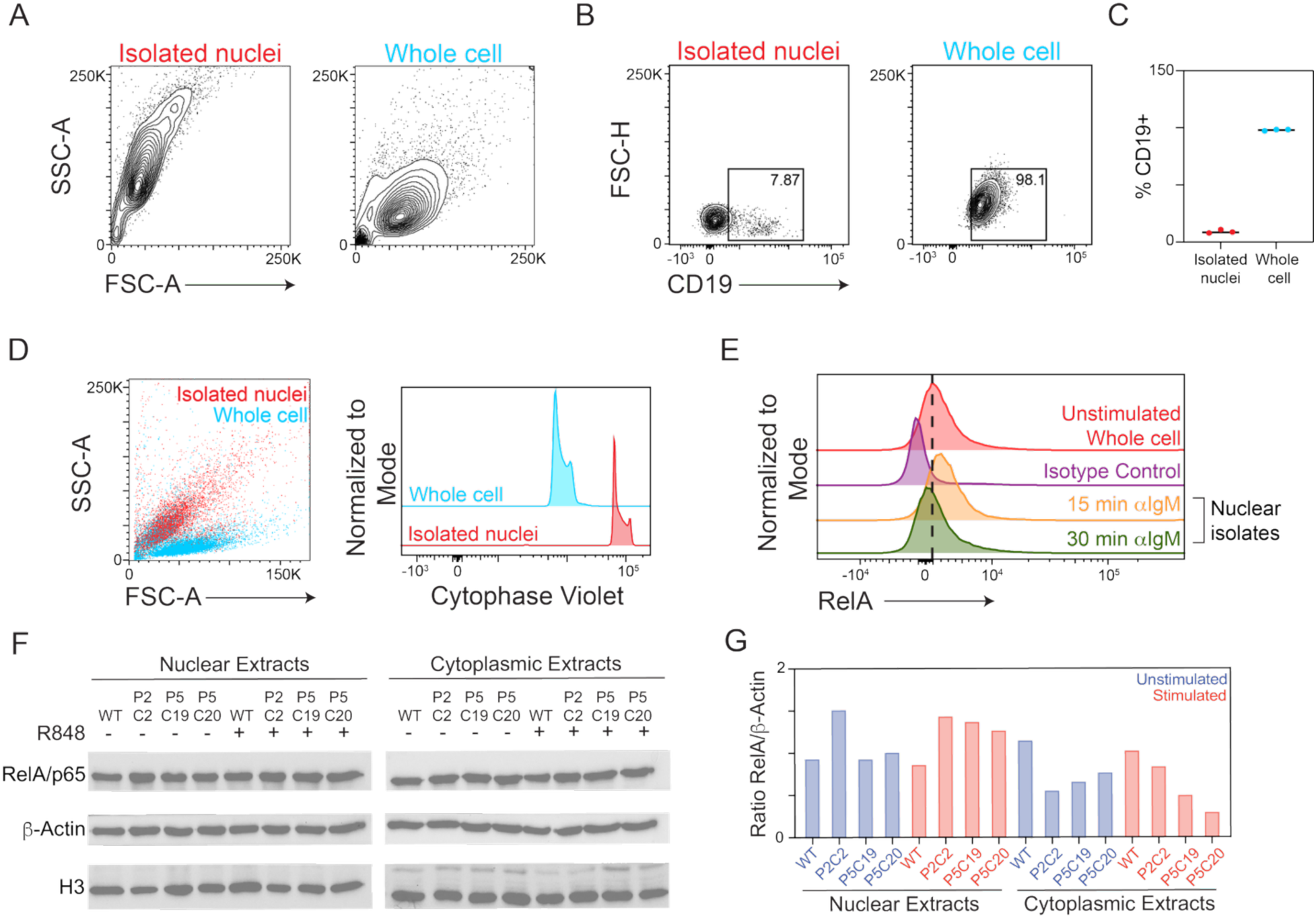
Insufficient RelA/p65 nuclear localization in stimulated Raji B cells. **A.** Forward and side scatter profiles of isolated nuclei (red) and whole cells (blue) by flow cytometry. **B.** Representative flow cytometry plots showing CD19 expression in isolated nuclei and whole cells, with percent CD19^+^ indicated. **C.** Summary quantification of percent CD19^+^ across isolated nuclei and whole cell samples across biological replicates (n=3). **D.** Overlay scatter plot of isolated nuclei (red) and whole cells (blue) showing distinct size populations (left), and representative histograms of Cytophase Violet DNA-binding dye intensity in isolated nuclei versus whole cells (right). **E.** Intracellular flow cytometry histograms of RelA/p65 expression in unstimulated whole cells, isotype control, and nuclear isolates from WT Raji cells stimulated with anti-IgM for 15- or 30-min. Dashed line indicates the position of the unstimulated whole cell peak. **F.** Western blot analysis of RelA/p65, β-actin, and histone H3 in nuclear and cytoplasmic fractions isolated from WT and A20-deficient Raji clones (P2C2, P5C19, P5C20) under unstimulated (-) and R848-stimulated (+) conditions. **G.** Quantification of RelA/p65 normalized to β-actin from nuclear and cytoplasmic fractions in **F** across unstimulated (blue) and stimulated (red) conditions.

Previous studies have also included DNA-binding dyes, such as propidium iodide, to confirm successful nuclear isolation relative to whole cells^62^. Consistent with this approach, we assessed DNA content using Cytophase Violet, a permanent DNA-binding dye (**Fig. 3D**). Isolated nuclei had a visibly higher expression of Cytophase violet compared to whole cells further confirming nuclear enrichment in our isolation protocol. Following validation of the isolation protocol, we next evaluated the nuclear localization of RelA/p65 as an indicator of NF-κB activation. Comparison of RelA/p65 in unstimulated whole cells to WT Raji cell nuclei stimulated with anti-IgM for either 15 or 30 min (**Fig. 3E**). The 15 min of anti-IgM stimulation showed a modest increase in RelA/p65 expression; however, the 30 min stimulation did not show a sustained increase in NF-κB activity. Overall, we expected a larger induction of RelA/p65 translocation to the nucleus following a robust anti-IgM stimulation. Although the approach of isolating nuclei was successful, we cannot confirm that there is nuclear accumulation of RelA/p65 in Raji B cells.

Furthermore, we wanted to confirm this phenomenon by examining both nuclear and cytoplasmic extracts from both WT and A20-deficient Raji cells. We performed western blot analysis on either WT Raji cells or A20-deficient clones (P2C2, P5C19, P5C20) and examined the presence of RelA/p65, β-actin, and histone H3 following stimulation with R848 (**Fig3. F-G**). RelA/p65 was detectable in both nuclear and cytoplasmic fractions; however, R848 stimulation did not produce a consistent increase in RelA/p65 across WT or A20 KO cells. Additionally, histone H3 and β-actin, which are normally confined to the nucleus and cytoplasm respectively, were present across all fractions. This phenomenon has been reported previously and does not necessarily mean that our nuclear and cytoplasmic fractions were not pure^62,63^. Together, these findings demonstrate that while nuclear isolation was successfully achieved, RelA/p65 nuclear translocation was not consistently inducible in Raji cells following stimulation with either anti-IgM or R848. These results suggest that RelA/p65 localization is not a reliable readout of NF-κB activation in this system under the conditions tested.

### Comparable NF-κB target gene expression in WT and A20-deficient B cells following TLR and BCR stimulation

Having observed no NF-κB activation, through either the phosphorylation of IκBα or nuclear accumulation of RelA in at the protein level, we next asked whether these differences were reflected in downstream transcriptional outputs of NF-κB signaling. We examined the mRNA expression of a panel of NF-κB target genes by quantitative PCR (qPCR) across a 24-hr time course following stimulation with either the R848 or anti-IgM (**Fig. 4A-B**). The transcriptional targets of NF-κB following B cell stimulation have been systematically characterized in murine B cells ^64,65^. We utilized panels of genes reported in these studies, including NF-κB subunits and regulators (*RELA, REL, TNFAIP3*), transcription factors induced downstream of NF-κB activation (*MYC, BHLHE40, IRF7, ATF4*), and an oxidoreductase with roles in redox regulation (*TXN*). Overall, the transcriptional responses were broadly comparable between WT and A20-deficient (P2C2) cells, with selective differences observed for specific genes under differing stimulation conditions.

**Figure 4.**
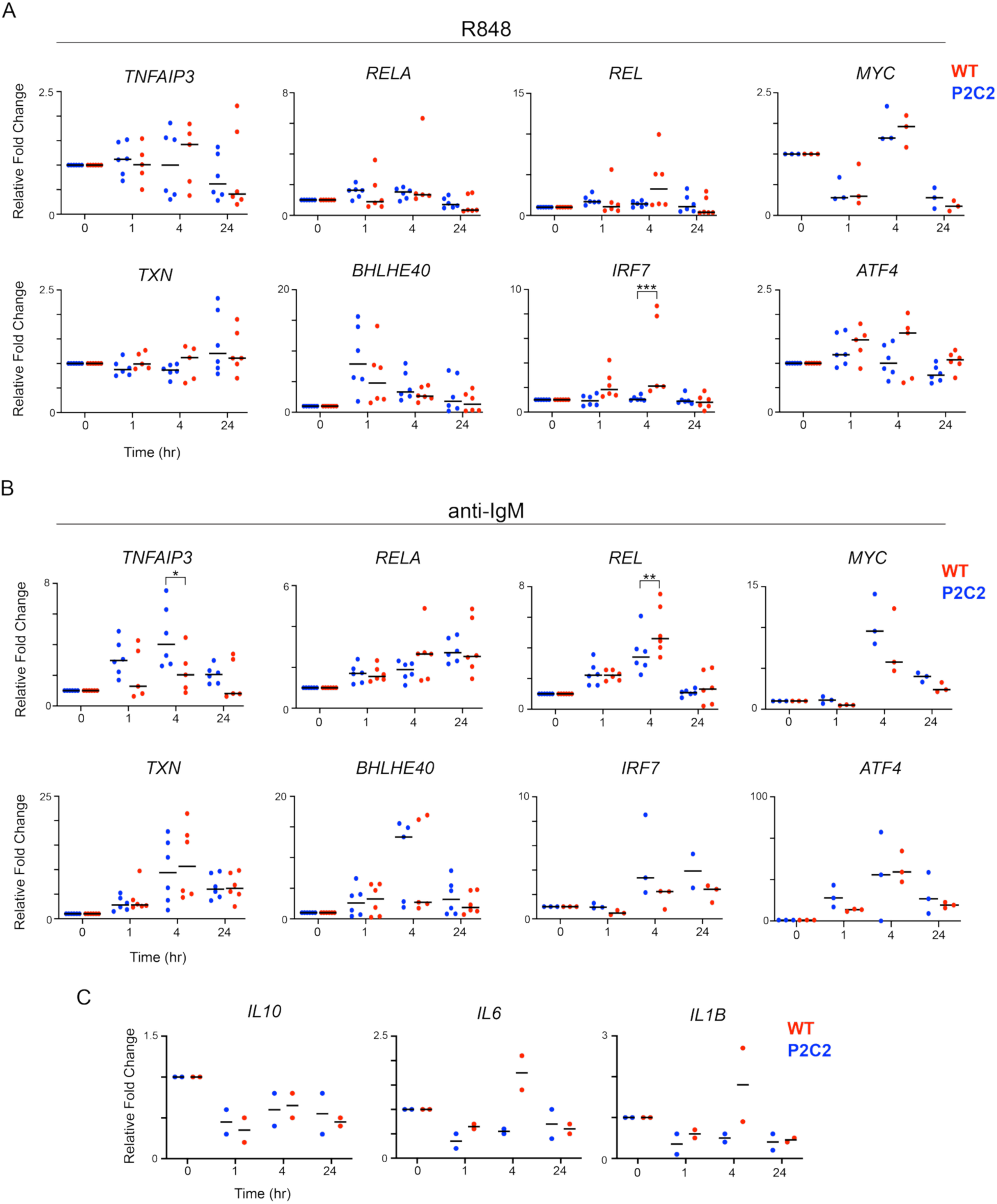
Comparable NF-κB target gene expression in WT and A20-deficient Raji cells following TLR and BCR stimulation. **A.** Relative fold change in mRNA expression of NF-κB target genes including NF-κB subunits and regulators (*TNFAIP3, RELA, REL*), downstream transcription factors (*MYC, BHLHE40, IRF7, ATF4*), and redox regulator (*TXN*). WT Raji cells (red) and A20-deficient clone P2C2 (blue) were evaluated over a 24-hr time course following R848 stimulation. Each dot represents an individual biological replicate and horizonal black bars indicate the median. Statistical significance is indicated. **B.** Relative fold change in mRNA expression of the same NF-κB target gene panel in WT and P2C2 Raji cells over a 24-hour time course following anti-IgM stimulation. Each dot represents an individual biological replicates, and color coordination is the same as **A**; horizontal bars indicate the median. Statistical significance is indicated. **C.** Relative fold change in mRNA expression of NF-κB-target cytokines *IL10*, *IL6*, and *IL1B* over a 24-hr time course following R848 stimulation. Each dot represents an individual biological replicate with WT (red) and A20-deficient cells (blue); horizontal bars indicate the median. All gene expression was normalized to a housekeeping gene (*18S*) and calculated relative to the unstimulated timepoint (0 hr). Statisical analysis was performed using an Ordinary two-way ANOVA with Tukey’s multiple comparisons test and single pooled variance.

Both R848 and anti-IgM stimulation induced *TNFAIP3* gene expression in the WT and P2C2. Although *TNFAIP3* mRNA signal was detectable in P2C2 cells by qPCR, this likely reflects amplification of a residual transcript fragment upstream or downstream of the CRISPR cut site, consistent with the absence of *TNFAIP3* protein confirmed by western blot (**Fig. 1**). Following R848 stimulation, *IRF7* expression was significantly increased in A20-deficient cells at 4-hr compared to WT, suggesting that the loss of *TNFAIP3*-mediated negative feedback may amplify *IRF7* induction downstream of TLR7/8 signaling. In contrast, under anti-IgM stimulation, *TNFAIP3* itself was significant upregulated in WT cells at 4-hr. An increase in *TNFAIP3* is consistent with its known role as a negative feedback regulator induced by NF-κB activation. Additionally, *REL* was elevated in P2C2 fold at 4 hr following anti-IgM stimulation, although there was considerable variability across biological replicates.

To assess the impact of *TNFAIP3* loss on cytokine production in human B cells, the expression of *IL10, IL6,* and *IL1β* was examined following R848 stimulation over a 24-hr time course (**Fig. 4C**). All three cytokines showed a declining expression pattern over 24-hr in both WT and P2C2 cells. However, there were no statistically significant differences observed between WT and P2C2. This suggests that in Raji cells, cytokine transcription is not overtly dysregulated by the loss of *TNFAIP3* following TLR7/8 stimulation. Together, these results demonstrate that loss of *TNFAIP3* does not broadly alter NF-κB-driven transcriptional responses following TLR7/8 or BCR stimulation, though selective differences in specific target genes such as *IRF7* and *REL* suggest that *TNFAIP3* may fine-tune rather than globally regulated NF-κB transcriptional outputs in Raji cells.

### Evaluation of A20 expression and NF-κB activation across lymphoblastoid B cell lines following TLR stimulation

Considering that Raji cells are derived from an individual with Burkitt lymphoma rather than representing a conventionally Epstein Barr virus (EBV)-immortalized lymphoblastoid cell line (LCL), we next wanted to determine whether stimulus-induced NF-κB activation could be more reliably detected in established LCLs. To determine if R848-induced NF-κB activation could be detected in an alternative B cell model system, we evaluated three EBV-immortalized LCLs including GM12878, GM12851, and GM12155. Each were stimulated with increasing concentrations of R848 (0-50μg/mL) for 15 or 30 min. A20 expression and NF-κB activity, assessed by phosphorylation and total levels of IκBα normalized to β-actin, were evaluated by western blot across all three lines (**Fig. 5A-F**).

**Figure 5.**
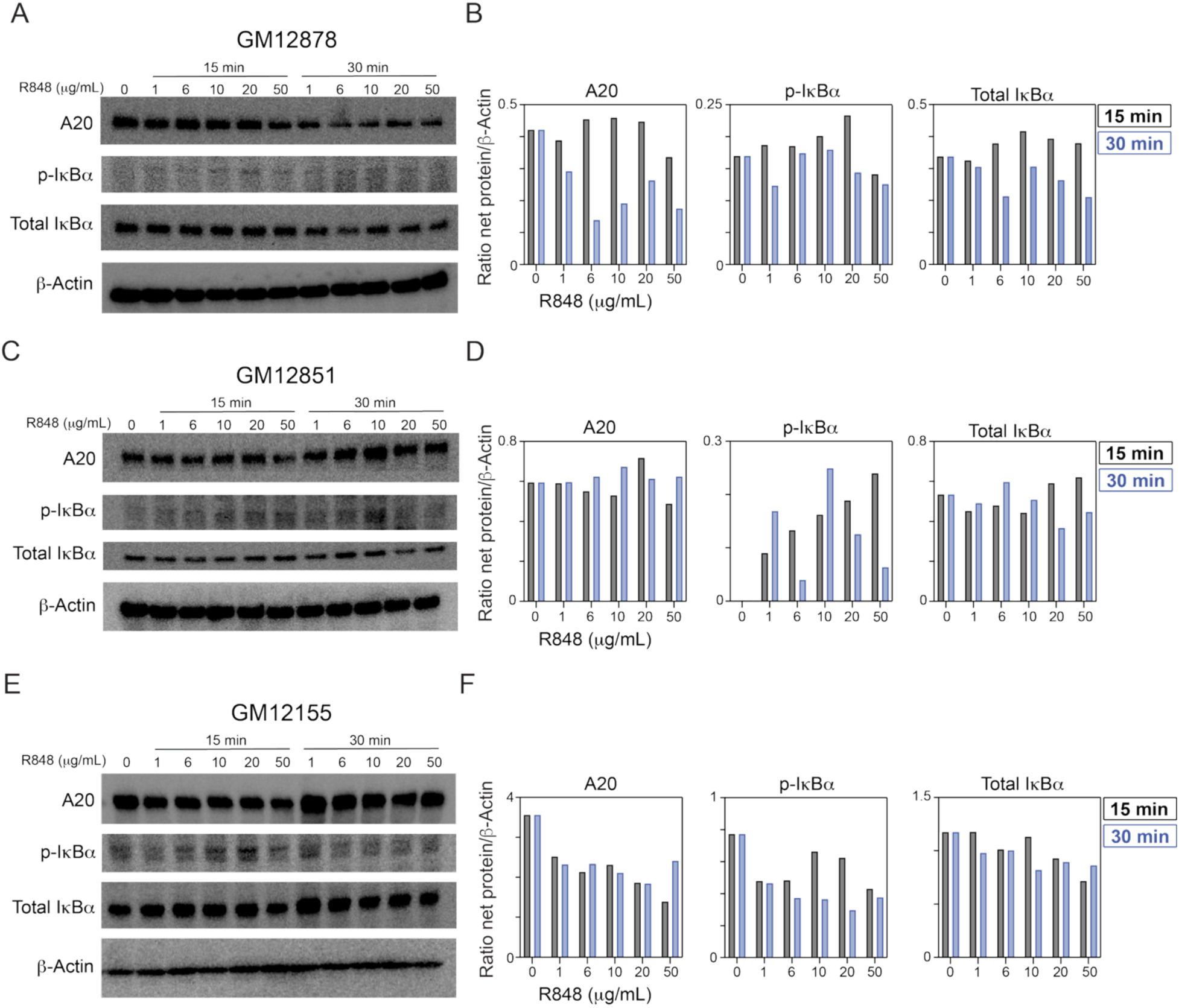
Assessment of A20 expression and canonical NF-κB activation across LCLs following R848 stimulation. **A.** Western blot analysis of A20, p-IκBα, total IκBα, and β-actin in GM12878 cells treated with increasing concentrations of R848 (0-50 µg/mL) at 15- and 30-min. **B.** Quantification of A20, p-IκBα, and total IκBα normalized to β-actin from **A** at 15 min (grey) and 30 min (blue). **C.** Western blot analysis of A20, p-IκBα, total IκBα, and β-actin in GM12851 cells treated with increasing concentrations of R848 (0-50 µg/mL) at 15- and 30-min. **D.** Quantification of A20, p-IκBα, and total IκBα normalized to β-actin from **C** at 15 min (grey) and 30 min (blue). **E.** Western blot analysis of A20, p-IκBα, total IκBα, and β-actin in GM12155 cells treated with increasing concentrations of R848 (0-50 µg/mL) at 15- and 30-min. **F.** Quantification of A20, p-IκBα, and total IκBα normalized to β-actin from **E** at 15 min (grey) and 30 min (blue).

A20 protein expression was constitutively expressed in all three GM LCLs at baseline and did not show consistent dose-dependent induction following R848 stimulation at either time point. Similarly, p-IκBα levels did not demonstrate a clear or reproducible pattern of induction across R848 dose range across any GM LCL cell line. Additionally, total IκBα was broadly stable in its expression and did not show reduction indicating NF-κB activation. Quantification of band intensities to the b-actin control confirmed the absence of stimulus dependent response across all three cell lines (**Fig. 5B, D, F**). Together, these findings indicate that R848 does not induce NF-κB activity in EBV-transformed LCLs under the conditions tested. This likely reflects constitutive NF-κB activation that limits stimulus-induced changes in culture. Consequently, these lines were not taken forward for further experimental use.

### Human primary B cells are a promising model for evaluating A20 function

Unexpectedly, the human B cell lines Raji and LCLs, failed to demonstrate robust NF-κB activation following A20 disruption. Given the known alterations in signaling pathways associated with viral transformation^66^, we hypothesized that these models may not accurately reflect endogenous NF-κB regulation. To address this, we turned to primary human B cells isolated from peripheral blood samples. To determine whether primary human B cells exhibit inducible NF-κB signaling, we first assessed the presence of A20 in both unstimulated and stimulated conditions, including diverse agonists of NF-κB such as TNF-α, anti-IgM, and R848 (**Fig. 6A**). Unstimulated primary human B cells showed low levels of A20 expression, while stimulation with TNF-α, anti-IgM, and R848 resulted in increased A20 protein levels. Visualization of the protein bands revealed that R848 stimulation induced the highest expression of A20 compared to both TNF-α and anti-IgM. As previously discussed, increased levels of A20 may indicate the induction of NF-κB through the positive feedback loop following signal transduction. To confirm NF-κB activation, we utilized qPCR analysis and assessed mRNA expression levels of NF-κB-target genes including *TNFAIP3, TNF, IL10,* and *IL6* during a 24-hr time course following R848 stimulation (**Fig. 6B**). Collectively, this analysis revealed a rapid induction of *TNFAIP3, TNF, IL10,* and *IL6* transcripts, peaking at 1-hr post stimulation and decreasing through 24-hr. Although all genes showed an induction, *IL10* and *IL6* had the highest expression at 1-hr with an almost 30-and 60-fold increase, respectively. Together, these results demonstrate that stimulation of human primary B cells induces A20 protein expression and upregulates NF-κB target genes, confirming that NF-κB is functionally activated in this model system.

**Figure 6.**
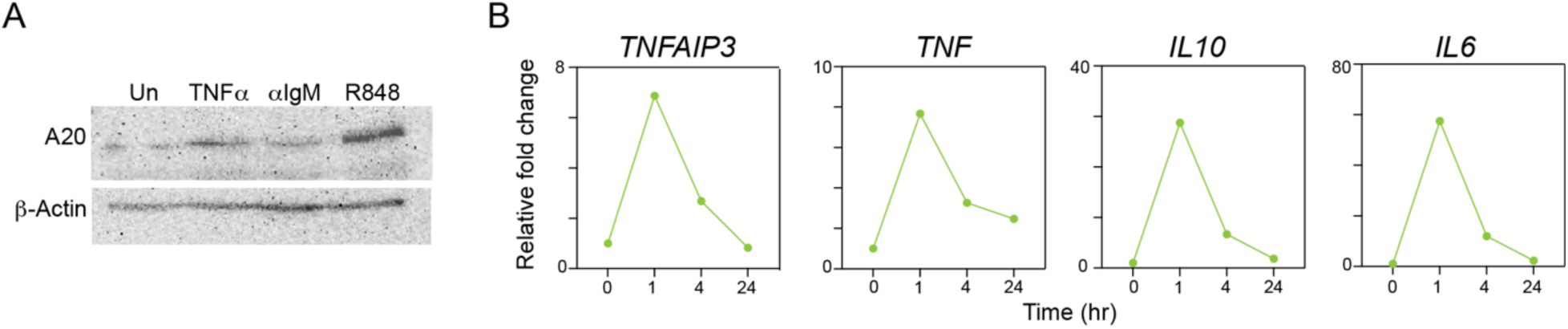
Primary human B cells represent an optimal model system for studying A20-dependent NF-κB regulation. **A**. Western blot analysis of A20 protein expression in primary human B cells under unstimulated conditions or following stimulation with TNF-α, anti-IgM, or R848, with β-actin as a loading control. **B.** Relative fold change in mRNA expression of NF-κB target genes *TNFAIP3, TNF, IL10,* and *IL6* in primary B cells over a 24-hr time course following R848 stimulation. All gene expression was normalized to a housekeeping gene (*18S*) and calculated relative to the unstimulated timepoint (0 hr).

## Discussion

A20 (*TNFIAP3*) is a critical negative regulator of NF-κB signaling and is essential for maintaining immune homeostasis across diverse cell types, as complete loss of A20 results in severe systemic inflammation and early lethality in mice^5^. Unlike other negative regulators of NF-κB, A20 is uniquely positioned to modulate pathway activation at multiple levels through its dual capacity as both a DUB and an E3 ubiquitin ligase, enabling coordinate control of the ubiquitination machinery that governs NF-κB signal transduction. Genetic variants and loss-of-function mutations in *TNFAIP3* have been associated with dysregulated inflammatory responses and the pathogenesis of autoimmune diseases, such as SLE and RA, highlighting the importance of tightly controlled NF-κB signaling in immune regulation^9,41,67^. As previously discussed, A20 function has been extensively characterized following TNF-α stimulation in HEK293, other cell lines, and murine models. Thus far, its functional role in human B cells remains incompletely understood. Because NF-κB occupies a central node in B cell activation, function, and fate determination, understanding how A20 shapes this pathway in a cell type-specific context is fundamental to defining its role in B cell-mediated autoimmunity. Therefore, we aimed to address this gap by pursuing a human B cell model system in which A20-dependent NF-κB regulation could be interrogated.

Our study successfully generated CRISPR/Cas9-mediated A20 knockout Raji B cells which we confirmed at the protein level by intracellular flow cytometry and western blot, and at the genomic level by Sanger sequencing and ICE analysis. However, despite robust validation of A20 loss, both WT and A20-deficient Raji cells failed to demonstrate inducible canonical NF-κB activation following TLR7/8 (R848) or BCR (anti-IgM) stimulation, as assessed by IκBα phosphorylation and degradation, RelA/p65 nuclear translocation, and NF-κB target gene expression. Raji cells are derived from a Burkitt lymphoma biopsy harboring a c-Myc translocation and carry latent EBV infection, both of which are known to constitutively engage oncogenic and inflammatory signaling networks^68,69^. Elevated baseline NF-κB signaling in Raji cells likely reduces the dynamic range required to observe additional activation following stimulation or A20 loss. Extending our study to three EBV-immortalized LCLs (GM12878, GM12851, GM12155) yielded a similar result, with no additional cell types demonstrating inducible IκBα phosphorylation or degradation following R848 stimulation. Importantly, EBV transformation drives constitutive NF-κB activation through LMP1 and LMP2A, which functionally mimic activated CD40 and BCR signaling, respectively^70–72^. This chronic mimicked signaling likely sustains NF-κB activity near its maximal threshold, thereby limiting the capacity for further pathway enhancement following stimulation with increasing dosages. Together, these findings demonstrate that viral transformation compromises the regulatory landscape of NF-κB in these models and highlights an important limitation of using transformed B cell lines for studying negative regulators, such as A20.

In contrast, primary human B cells isolated from peripheral blood demonstrated robust and inducible NF-κB activation, establishing them as the more appropriate model system for this work. Stimulation with R848 induced clear upregulation of A20 protein and qPCR analysis confirmed rapid transcriptional induction of NF-κB target genes including *TNFAIP3, TNF, IL6*, and *IL10*. This kinetic profile is consistent with canonical NF-κB activation followed by negative feedback-mediated resolution, and closely mirrors patterns previously reported in other primary immune cell types^64,65^. Together, these findings support that primary human B cells exhibited a responsiveness to NF-κB stimulation and allow for a clear detection of experimental readouts that could not be achieved in Raji or LCLs.

Collectively, this work makes several important contributions by demonstrating that transformed B cell lines, including Raji and LCLs, are not suitable models for studying A20 dysregulation following NF-κB activation. This was a surprising result given their broad usage historically in literature. Primary human B cells are a usable system in which NF-κB is robustly induced and A20 expression can be evaluated; however, tools to robustly edit and manipulate primary B cells are largely lacking from the literature. Together, these findings establish the experimental framework upon which subsequent studies may be built. Most importantly, they underscore that cellular contexts and stimulation conditions are critical variables that must be carefully considered when studyingA20 and NF-κB biology in human B cells.

## Methods

### Cell lines and culture conditions

Raji B cells were cultured in RPMI-1640 supplemented with 10% fetal bovine serum (FBS) and 1% penicillin-streptomycin at 37°C with 5% CO₂. LCLs GM12878, GM12851, and GM12155 were maintained under equivalent RPMI-based conditions. Human embryonic kidney 293FT (293FT) cells were cultured in DMEM supplemented with 10% FBS. Cell lines were obtained through ATCC or the Coriell Institute for Medical Research.

### CRISPR/Cas9-mediated knockout of TNFAIP3 in Raji cells

Four single guide RNAs (gRNAs) targeting the TNFAIP3 locus (gRNA #1, #2, #5, #6) and Cas9 were nucleofected into Raji cells using the SG Cell Line 4D-Nucleofector™ X Kit (Lonza, V4XC-3012). Edited cells were surface stained with a fixable viability dye (BioLegend, 423113) prior to fixation/permeabilization (Thermofisher, 88-8824-00). Intracellular staining was then performed to assess A20 protein loss using anti-human polyclonal A20 antibody (ThermoFisher, 23456-1-AP). A corresponding isotype control (Biolegend, 405314) antibody was used to assess nonspecific binding and establish background fluorescence. Data was collected using BD FACSymphony. To generate clonal knockout lines, parental pools transduced with gRNA #2 and #5 were subjected to single-cell sorting using a sequential gating strategy (FSC-A/SSC-A, FSC-A/FSC-H, Live/Dead) using BD FACSAria II. All flow cytometry data was analyzed using Flowjo v.10. Expanded single-cell clones were screened by intracellular flow cytometry and mean fluorescence intensity (MFI) quantification.

### ICE Analysis

Genomic disruption at the *TNFAIP3* locus was confirmed in the P2C2 clone by isolating genomic DNA (Zymo, D3021). Targeted region surrounding the cut site was amplified by PCR using primer set v3/v4 and purified (Thermo Scientific, FERK0701). PCR amplicons were Sanger sequenced, with indel analysis performed using the Inference of CRISPR Edits (ICE) algorithm.

### Cell stimulation

Raji cells and LCLs were stimulated with the TLR7/8 dual agonist Resiquimod (R848) (Invivogen, tlrl-r848) at concentrations ranging from 0–50 µg/mL for the indicated timepoints. B cell receptor (BCR) stimulation was performed using anti-IgM (Jacksonimmuno, 109-006-129). 293FT cells were stimulated with recombinant TNF-α (ThermoFisher, A42552) to validate NF-κB signaling reagents.

### Western blot analysis

Whole-cell lysates were prepared from stimulated or unstimulated cells and resolved by SDS-PAGE. Proteins were transferred to membranes and probed with antibodies against A20 (Cell Signaling, clone D13H3, #5630) phospho-IκBα (p-IκBα) (Cell Signaling, 2859T), total IκBα (Cell Signaling, 4814T), RelA/p65 (Cell Signaling, 8242T), histone H3 (Cell Signaling, 4499T), and β-actin (loading control) (Santa Cruz, sc-47778). Bands were quantified by densitometry (ImageJ 1.54g) and normalized to β-actin.

### Nuclear and cytoplasmic fractionation

Nuclear and cytoplasmic fractions were isolated using an extraction kit (ThermoFisher, 78833) from both WT and A20-deficient Raji clones (P2C2, P5C19, P5C20) under unstimulated and R848-stimulated conditions. Nuclear purity was assessed by flow cytometry using CD19 (Biolegend, 363010) surface staining and Cytophase Violet DNA-binding dye (Biolegend, 425701), and by western blot using histone H3 (nuclear marker) and β-actin (cytoplasmic marker). RelA/p65 localization across fractions was evaluated by western blot.

### Intracellular RelA/p65 flow cytometry

A20 protein expression was assessed by intracellular staining as previously described. RelA/p65 nuclear localization was evaluated by intracellular staining in isolated nuclei and whole cells following a previously published protocol^62^. A coordinating isotype control was also included, and both were stained with a conjugated secondary antibody. MFI was calculated for each condition as previously described.

### RNA isolation and quantitative PCR (qPCR)

Total RNA was extracted (Zymo, R1054) from stimulated cells at the indicated timepoints. cDNA was synthesized and gene expression was assessed by qPCR for NF-κB target genes including *TNFAIP3, RELA, REL, MYC, BHLHE40, IRF7, ATF4, TXN, IL10, IL6,* and *IL1B*. All primer sequences are found in **Table 1**. All expression was normalized to the housekeeping gene 18S and expressed as fold change relative to the unstimulated (0 hr) timepoint. Statistical analysis was performed using an Ordinary two-way ANOVA with Tukey’s multiple comparisons test, with single pooled variance.

**Table 1.**
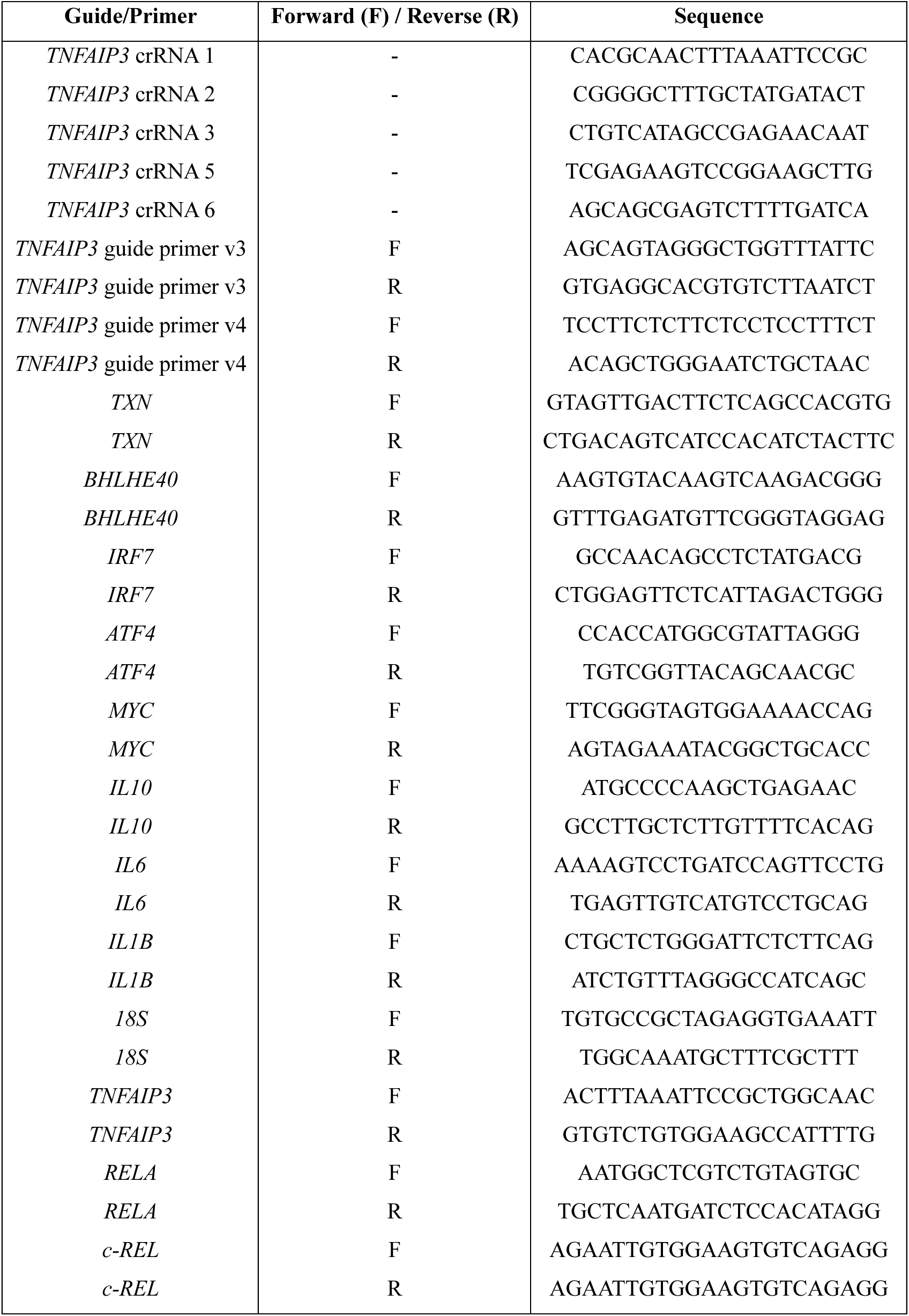
Guide and Primer Sequences.

### Primary human B cell isolation and culture

Peripheral blood B cells were isolated from fresh or cryopreserved PBMCs by negative selection (StemCell, 17954). TLR stimulation was performed with R848 (TLR7/8) or ODN2006 (TLR9 agonist) (Invivogen, tlrl-2006-1) in IMDM (ThermoFisher, 12440061) media.

### Data visualization

Graphs and data visualizations for flow cytometry, qPCR, and western blot experiments were generated using GraphPad Prism (v10.5.0) and Microsoft Excel (v16.102.1).

## Acknowledgements

The authors thank the members of the Scharer and Boss labs for their critical discussions and insights as well as the Emory Integrated Flow Cytometry Core.

## Author Contributions

R.K. contributed to investigation, formal analysis, writing original draft, and visualization. C.D.S. contributed conceptualization, supervision, writing review, editing, and funding acquisition.

